# Statistical Causal Discovery in Developing and Refining Adverse Outcome Pathway (AOP)

**DOI:** 10.1101/2025.08.31.672289

**Authors:** Kyoshiro Hiki, Thong Pham, Michio Yamamoto, Takehiko I. Hayashi, Shohei Shimizu

## Abstract

Statistical causal discovery (SCD) has the potential to advance the development and evaluation of Adverse Outcome Pathways (AOPs) by inferring causal relationships directly from data. However, ecotoxicology data often has challenges for SCD applications, such as missing data and violation of SCD algorithm assumptions. As a proof-of-concept, we applied a linear non-Gaussian acyclic model (LiNGAM), a representative SCD method, to three types of ecotoxicology datasets: (1) bivariate dose–response relationships, (2) bivariate response– response relationships, and (3) a multivariate dataset with a known causal structure. Missing data were addressed through multiple imputation followed by causal estimation using DirectLiNGAM, a direct method for estimating LiNGAM. DirectLiNGAM identified correct causal directions with high statistical reliabilities in three of four bivariate dose-response cases, even when assumptions such as linearity and non-Gaussianity were partially violated. In contrast, response–response cases did not yield a single dominant direction, likely due to the limited number of replicates. In the multivariate case, the inferred graphs closely resembled the expert-curated causal graph, achieving high recall (0.50–0.75), despite relatively low precision (0.31–0.40). These results demonstrate the utility of SCD, combined with multiple imputation, in identifying relevant key events, revealing missing links, and refining existing AOP and quantitative AOP (qAOP) models, under realistic ecotoxicological constraints.

**Synopsis:** Statistical causal discovery can advance the development of adverse outcome pathways in a data-driven manner, enabling efficient chemical risk assessment.

**TOC Art:** 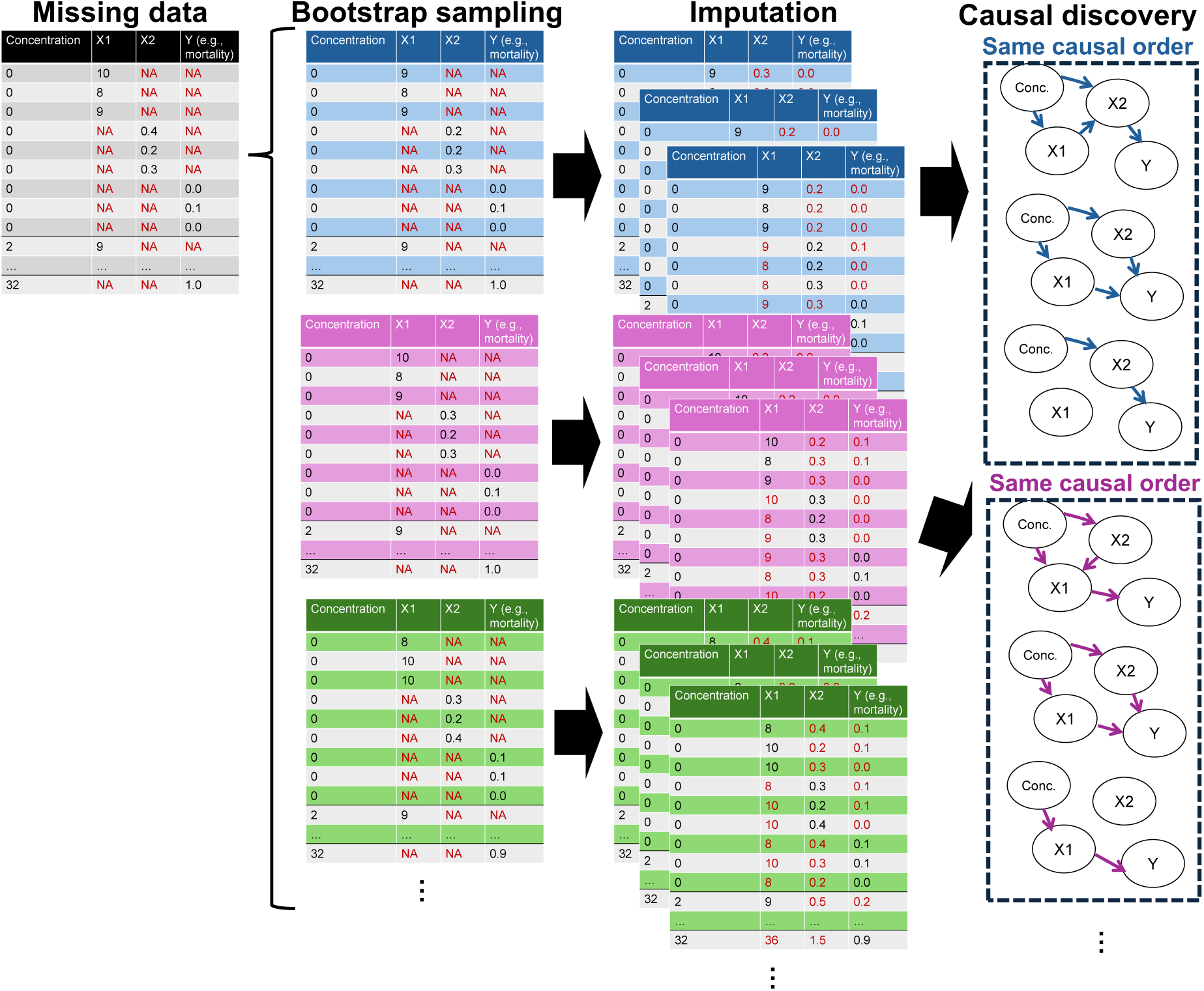

## 1. INTRODUCTION

Rapid advances in in vitro testing, in silico modelling, and high-throughput omics technologies have flooded environmental toxicology with rich but complex datasets. Making sense of these data requires frameworks that explicitly connect molecular-scale perturbations to ecologically relevant impacts. The adverse outcome pathway (AOP) framework has evolved as a promising tool for this purpose, providing a structured representation of the biological key events (KEs) that link a molecular initiating event (MIE) to adverse outcomes (AO) at higher levels of biological organization (e.g., death, reproduction impairment) (Ankley et al. 2010). Most published AOPs are still developed by expert judgment, although several semi-automated techniques have recently been proposed, such as text mining (Carvaillo et al. 2019; Corradi et al. 2022) and large language models (LLMs) (Shi and Zhao 2024). Consequently, KE selection and causal directions are often inferred manually, and the resulting networks can be incomplete and subjective (Perkins et al. 2015; Spînu et al. 2022; Jeong et al. 2025; Zhou et al. 2025). As the volume of mechanistic toxicology data grows and the number of chemicals increases, there is an urgent need for data-driven approaches that can (i) propose new AOP structures and (ii) verify and refine existing ones.

Statistical causal-discovery (SCD) methods offer such a data-driven solution. These algorithms infer causal structure directly from observational and/or interventional data, by imposing tractable assumptions that restrict the search space of possible causal graphs (Shimizu and Blöbaum 2020). Traditional SCD methods, such as Peter and Clark (PC) and fast causal inference (FCI) algorithms (Spirtes et al. 2001), do not necessarily identify a fully directed graph and often leave several directions unresolved. In other words, the output is a set of causal structures satisfying the same conditional independence (i.e., Markov-equivalence class). However, some advanced SCD methods can distinguish among causal structures within the same Markov-equivalence class, by making additional assumptions. For example, in the bivariate case, the linear non-Gaussian acyclic model (LiNGAM) distinguishes X → Y from Y → X when noise terms are non-Gaussian and independent (Shimizu et al. 2006). SCD has been applied to a wide range of research fields, such as neurobiology (Chiyohara et al. 2023), molecular biology (Maathuis et al. 2010; Feng et al. 2023), epidemiology (Rosenström et al. 2012; Lopez Barrera and Miljkovic 2022; Ribeiro-Dantas et al. 2024), and industrial engineering (Cao et al. 2022). Nevertheless, its use in environmental toxicology remains limited (Trairatphisan et al. 2021; Tamura et al. 2024). To our knowledge, no study has systematically integrated SCD into AOP development.

Integrating SCD with AOPs is conceptually attractive for four reasons. First, although chemical concentration or dose can be randomized, experimental randomization of upstream KEs is often infeasible (Zhou et al. 2025), making SCD a practical alternative for causal inference. Second, high-dimensional omics or phenotypic screening data can be searched for causal chains, revealing novel pathways not limited by expert pre-conceptions. Third, for partially built AOP networks, SCD can re-discover the existing edges and uncover missing or spurious links. Lastly, causal coefficients derived by SCD algorithms can be used to estimate response–response relationships between KEs, thereby extending efforts in quantitative AOP (qAOP) modelling.

However, there are several challenges in applying SCD to AOP development. AOP construction requires toxicity data across multiple levels of biological organization, but measurements at the molecular, cellular, or organ levels typically require destructive sampling. As a result, biological endpoints cannot be measured in the same biological replicate and are often measured in separate replicates, although exposure dose or concentration, a non-destructive variable, can be measured for all replicates. This results in datasets with missing values particularly for biological endpoints (Figure 2). Because there is currently no consensus on how to handle missing values in SCD (Glymour et al. 2019), it is necessary to establish practical approaches for managing missing data in the context of AOP development. Another challenge is that toxicology data often violate assumptions of SCD, such as linearity, acyclicity, and non-Gaussian error distributions. For example, toxicology data frequently exhibit sigmoidal (S-shaped) relationships and feedback loops, which conflict with common SCD assumptions. While one might attempt to verify these assumptions using statistical tests and apply SCD only to datasets that satisfy them, this approach is often impractical due to the limited statistical power (Prakash et al. 2024) inherent in toxicology datasets that typically have small sample size. Instead, there is a growing need for SCD methods that are robust to violations of underlying model assumptions (Montagna et al. 2023).

The present study represents the first systematic attempt to incorporate SCD into the AOP framework for environmental risk assessment. Our objectives are to: (i) evaluate whether SCD can reliably identify the causal direction in ecotoxicology data that often violate its underlying assumptions, using bivariate datasets with known causal directions; and (ii) assess the utility of SCD for developing and validating an AOP by comparing reconstructed graphs with the canonical pathway. For both objectives, to address missing values, we performed multiple imputation and then assessed the statistical reliability of inferred causal graphs based on the bootstrap probabilities. We used DirectLiNGAM (Shimizu et al. 2011), a direct method for estimating LiNGAM, as the SCD method. Finally, we discussed limitations and future directions for SCD-enabled AOP development.

## 2. Materials and Methods

### 2.1. Theory

DirectLiNGAM makes four assumptions: (i) relationship among variables is linear, (ii) the causal graph is acyclic, (iii) error terms are non-Gaussian, and (iv) there are no unobserved common causes (Shimizu et al. 2011). Suppose that we have observed data of *n* variables (i.e., *n* kinds of toxicity endpoints) and the data can be expressed as a vector ***X*** = (*x*_1_, *x*_2_, …, *x*_n_). Suppose that ***X*** is generated from a process represented graphically by a directed acyclic graph (DAG), analogous to AOP or AOP network. An example of DAG with three variables is shown in Figure 1A. The DAG can be expressed as a *n* × *n* adjacency matrix ***B***, where each component *b*_ij_ represents the connection strength from a variable *x*_j_ to *x*_i_ in the DAG (Figure 1C). Note that this indexing is the reverse of typical use in graph theory, where *b*_ij_ represents a link from *x*_i_ to *x*_j_. The structural causal model (Pearl 2009) of DirectLiNGAM can be given as follows (Figure 1B):

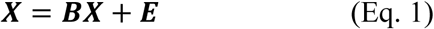

where ***E*** is a vector of error terms and the matrix ***B*** can be permuted to a strictly lower triangular matrix due to the assumption of acyclicity.

**Figure 1.**
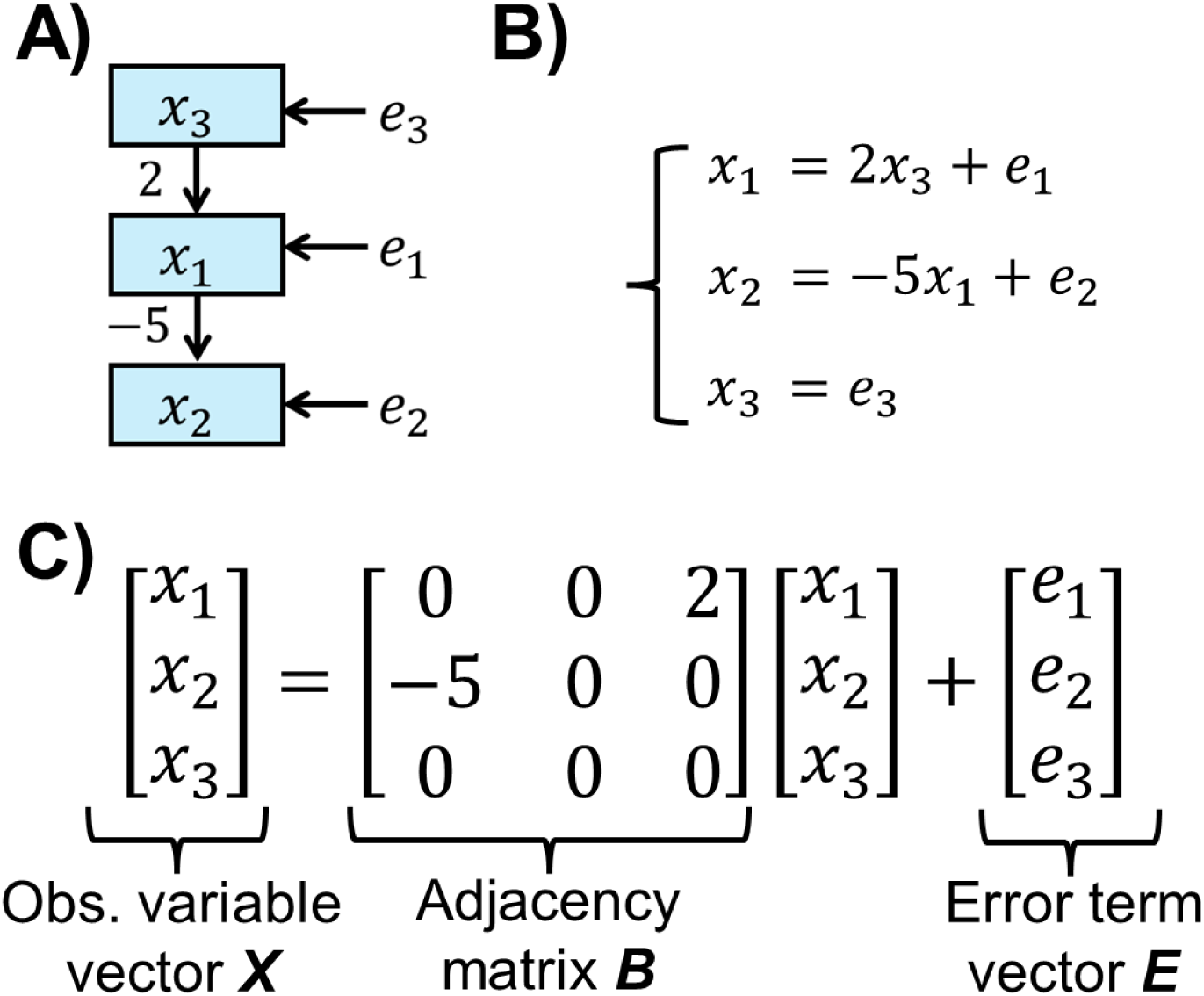
Illustration of the linear non-Gaussian acyclic model (LiNGAM) and directed-acyclic graph (DAG). The same causal relationship is illustrated in different forms: (A) DAG, (B) structural equation model, and (C) matrix model.

To estimate the matrix ***B*** in DirectLiNGAM, the algorithm identifies exogenous variables, defined as variables with no parents, through iterative pairwise regression. The algorithm first finds an exogenous variable (e.g., *x*_3_ in Figure 1) based on their independence from the residuals of pairwise regression with other variables. In practice, the exogenous variable is identified as the one most independent from its residuals, typically assessed using mutual information or other independence measures. Next, the algorithm identifies the “second” exogenous variable (e.g., *x*_1_ in Figure 1) in the causal ordering based on the independence of the first exogenous variable from the residuals of the remaining variables. Because the causal order of residuals preserves the ordering of the original variables, this recursive approach can be used to infer the full causal ordering. These processes are theoretically supported by the assumptions of DirectLiNGAM and two lemmas in Shimizu et al. (2011).

Although DirectLiNGAM can be run without any prior knowledge on causal structure, more efficient inference can be achieved if prior knowledge on partial structure is available. When we apply SCD to AOP development, stressor or chemical exposure should be exogenous variable and AO should be set as a sink variable, the one with no child variable. If an exogenous variable is identified based on prior knowledge, we can skip the step of evaluating independence between the observed variable and the residuals of other variables. Similarly, if a sink variable is known, we can exclude it from further independence assessments, as it does not serve as a parent in the causal graph. These processes can improve inference accuracy and reduce computational cost. Note that while AOP, by definition, begins with an MIE and is therefore chemical agnostic (Ankley et al. 2010), a stressor gradient may be sometimes incorporated into AOP analysis to estimate quantitative relationship between KEs (Moe et al. 2020). In addition, if causal relationships between KEs are known (e.g., upregulation of gene A directly causes expression of gene B), such knowledge can be incorporated as prior model input.

### 2.2. Case studies for applying SCD to AOP development

We applied DirectLiNGAM to ecotoxicology data with known causal directions and structure, with an aim to show that SCD methods can be useful to AOP development even in the presence of two challenges: missing data and violations of model assumptions. We present three ecotoxicology case studies: i) dose-response bivariate relationships, ii) response-response bivariate relationships, and iii) multivariate relationships. In the latter two cases, original data included missing values and then we addressed this by multiple random imputation. For all the three cases, the statistical reliability of inferred causal graphs was assessed based on the bootstrap probabilities (Komatsu et al. 2010). An example of the python code is available on GitHub (https://github.com/KyoHiki/SCDinAOP).

#### 2.2.1 Case study 1: bivariate dose-response relationship

Dose-response datasets were obtained from the drc R package ver. 3.0.1 (Ritz et al. 2015) and from a previous study (Cedergreen et al. 2016). We analyzed the causal direction between concentration of a test chemical and four endpoints: earthworm survival, mysid dry weight, the number of offspring, and fluorescence as a proxy of cell viability (Table 1). As each dataset originated from a single experiment in which concentration and the biological response were recorded simultaneously, there were no missing values. We ran DirectLiNGAM with 1000 bootstrap iterations using the “DirectLiNGAM” and “bootstrap” functions in the lingam Python package ver. 1.9.1 (Ikeuchi et al. 2023). The inferred causal directions were compared with the known directions, where chemical concentration is the parent and biological response is the child variable.

**Table 1.**
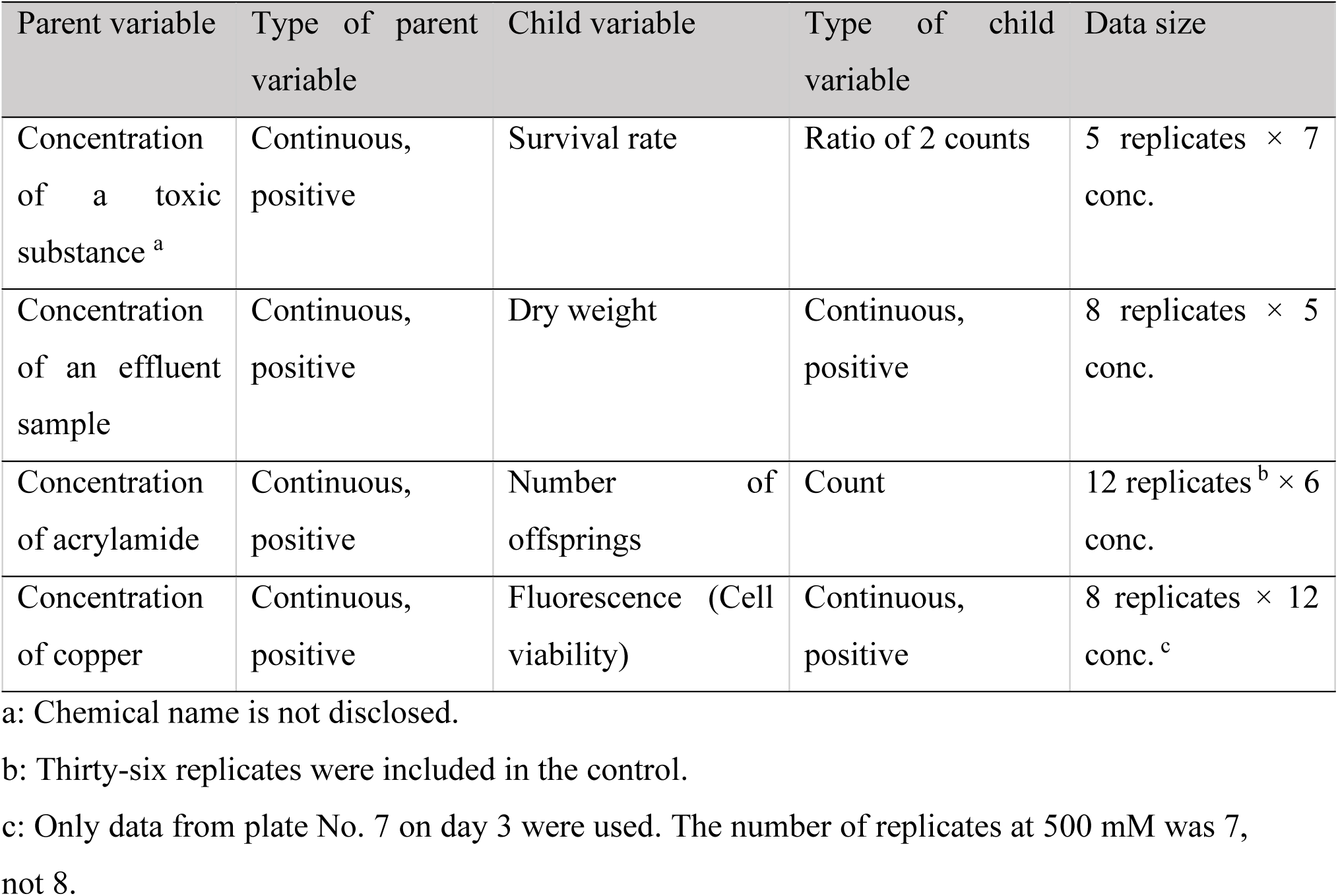
Bivariate dose-response ecotoxicology datasets with known causal direction.

| Parent variable | Type of parent variable | Child variable | Type of child variable | Data size |
| --- | --- | --- | --- | --- |
| Concentration of a toxic substance <sup>a</sup> | Continuous, positive | Survival rate | Ratio of 2 counts | 5 replicates × 7 conc. |
| Concentration of an effluent sample | Continuous, positive | Dry weight | Continuous, positive | 8 replicates × 5 conc. |
| Concentration of acrylamide | Continuous, positive | Number of offsprings | Count | 12 replicates <sup>b</sup> × 6 conc. |
| Concentration of copper | Continuous, positive | Fluorescence (Cell viability) | Continuous, positive | 8 replicates × 12 conc. <sup>c</sup> |

a: Chemical name is not disclosed.
b: Thirty-six replicates were included in the control.
c: Only data from plate No. 7 on day 3 were used. The number of replicates at 500 mM was 7, not 8.

To evaluate whether the data satisfied the assumptions of DirectLiNGAM, we applied three diagnostic statistical hypothesis tests. Linearity was assessed with the Ramsey Regression Equation Specification Error Test (RESET) (Ramsey 1969), which checks the significance of the quadratic term in an ordinary least-squares regression. The RESET was performed using the linear_reset function in the statsmodels Python package ver. 0.14.4 (Seabold and Perktold 2010) with the argument power = 2. Normality of the regression residuals was evaluated using the Shapiro–Wilk test. Finally, to evaluate the LiNGAM assumption that there are no unobserved common causes, we examined independence between error terms using the Hilbert–Schmidt Independence Criterion (HSIC) (Gretton et al. 2005), through the get_error_independence_p_values function of the lingam package (Ikeuchi et al. 2023). In addition to HSIC, the F-correlation was estimated as the nonlinear correlation between two variables (Bach and Jordan 2002), using the f_correlation function.

#### 2.2.2. Case study 2: bivariate response-response relationship

Response-response dataset (Table 2) was obtained from Moe et al. (2020), which reports the effects of 3,5-dichlorophenol (DCP) on the aquatic plant *Lemna minor* (original experiments by Xie et al. 2018). The measured endpoints are oxidative phosphorylation (OXPHOS), reactive oxygen species (ROS), electron transfer rate (ETR), maximum quantum yield of photosystem II (Fv/Fm), lipid peroxidation (LPO), and number of leaves (frond number) (Table 2). These represent the key events of an AOP from the MIE (reduction in OXPHOS and production ROS) to AO (reduced frond number). DCP concentrations were 0, 0.5, 1, 1.5, 2, 3, 4, and 8 mg/L; but the highest two concentrations were excluded from the analysis because most endpoints were not recorded due to high plant mortality.

**Table 2.** Bivariate response-response ecotoxicology datasets with known causal direction.

| Parent variable | Type of parent variable | Child variable | Type of child variable | Data size |
| --- | --- | --- | --- | --- |
| OXPHOS | Count | Frond number | Count | 3 replicates × 6 conc. |
| ETR | Continuous, positive | Frond number | Count | 3 replicates × 6 conc. |
| ROS | Count | Frond number | Count | 3 replicates × 6 conc. |
| Fv/Fm | Ratio of 2 counts | Frond number | Count | 3 replicates × 6 conc. |
| LPO | Continuous, positive | Frond number | Count | 3 replicates × 6 conc. |

OXPHOS: Oxidative phosphorylation, ETR: Electron transfer rate, Fv/Fm: Maximum quantum yield of photosystem II, LPO: Lipid peroxidation, ROS: Reactive oxygen species. Data were taken from Moe et al. (2020). Each variable was measured separately; therefore, we reproduced pseudo-replication by randomly imputing missing values, and then applied SCD method to each imputed dataset (see Figure 2).

As each variable was measured in separate biological replicates, initial data tables contained only concentration and a single response variable, with all other variables recorded as missing (Figure 2). To construct datasets suitable for SCD, we employed hot-deck multiple imputation (Rubin 1987), a non-parametric imputation technique. Each missing value was imputed by random resampling of observed values within the same concentration group (Figure 2). The imputation procedure was repeated 50 times for each bootstrap sample to generate an ensemble of fully imputed datasets, and DirectLiNGAM was applied to every imputed dataset for two variable combinations in Table 2, using the “MultiGroupDirectLiNGAM” function, under the assumption that causal orders are common across all imputed datasets. We then performed 500 iterations of bootstrap sampling, imputation, and DirectLiNGAM, and compared a total of 25,000 inferred causal directions with the known directions (Table 2), where frond number is the child and the other biological response is the parent variable.

**Figure 2.**
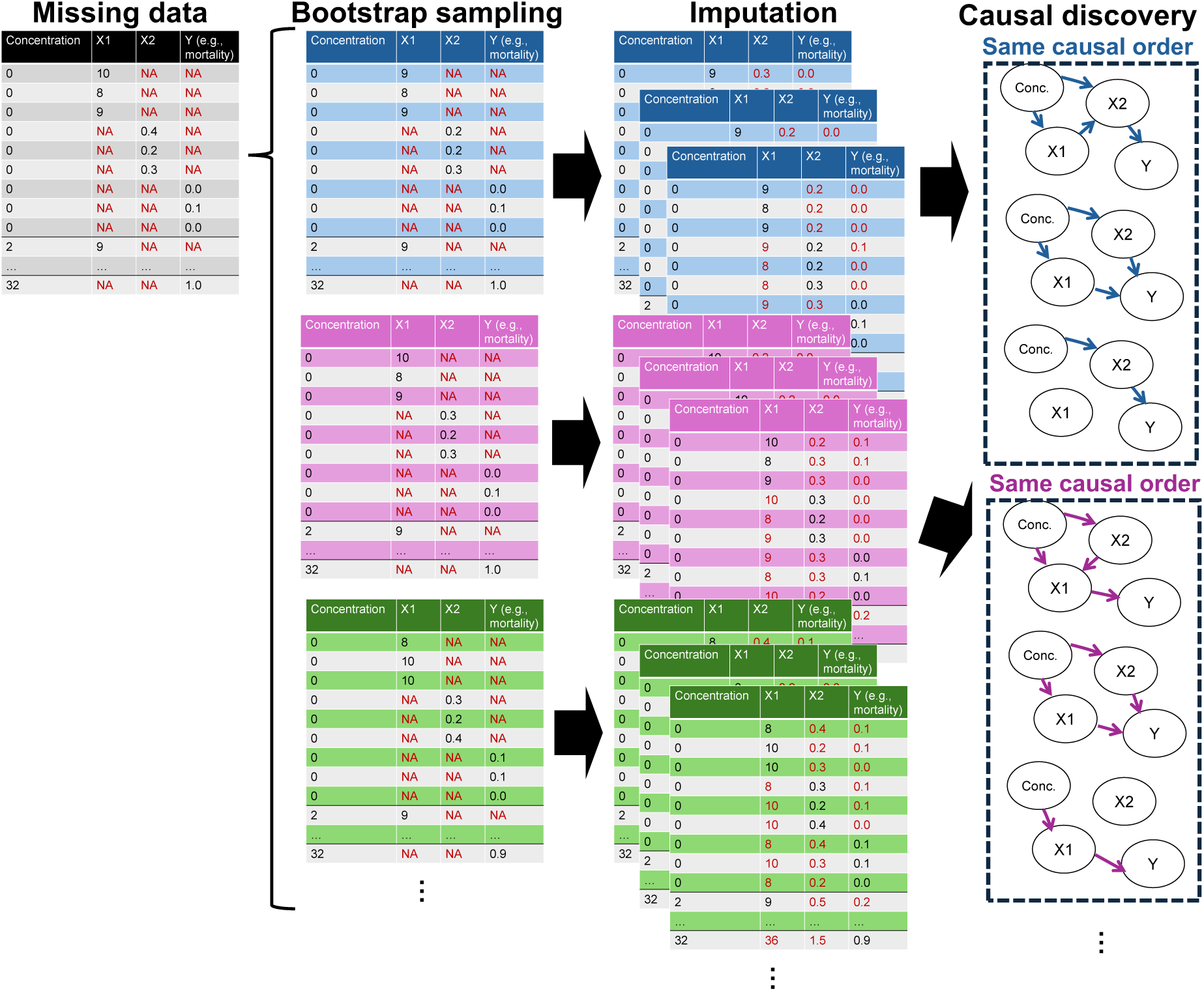
Workflow for applying statistical causal discovery (SCD) to infer an adverse outcome pathway (AOP) from a toxicity dataset that contains missing values. Original data have observed variables (X1, X2, and outcome Y) with missing values (NA) at each test concentration. For each bootstrap sample, hot-deck multiple imputation is performed, where missing values are randomly replaced with observed ones from the same concentration level. This imputation procedure is repeated 50 times to generate fully imputed datasets. An SCD algorithm is applied to each imputed dataset to recover a directed acyclic graph (DAG), assuming a common causal order across datasets from the same bootstrap sample. Finally, the probability of DAGs and paths is then estimated.

Based on the pattern of missingness in the data, it is reasonable to regard the missing-data mechanism as missing at random (MAR); under this condition, hot-deck multiple imputation is known to yield valid statistical inference (Reilly 1993). In addition, our preliminary investigation using simulated MAR data supported the use of hot-deck multiple imputation, showing that it yielded the most stable and accurate estimates of causal effects in linear relationships (Text S1 and Table S1).

#### 2.2.3. Case study 3: multivariate relationship

Beyond bivariate relationships, a multivariate dataset with a known causal structure was used to further test whether SCD methods can be useful in developing novel AOP network or in refining partially developed AOP network. The toxicity data of DCP on *L. minor* (Moe et al. 2020), same data as in the last section, was used also for multivariate case example. As with the last section, we generated 50 fully imputed datasets by random within-concentration resampling. Each imputed table contained seven variables: DCP concentration, OXPHOS, ETR, ROS, Fv/Fm, LPO, and frond number. DirectLiNGAM was applied to every dataset with 5000 bootstrap iterations, both with and without prior knowledge that (i) concentration is an exogenous variable and (ii) frond number is a sink (Figure 3C). The number of bootstrap iterations was increased compared to the last section (500 times), because the multivariate causal graph was more complex, which reduced the frequencies for individual DAGs. Increasing the number of bootstrap iterations allowed us to detect and report even less frequent graphs with greater reliability.

**Figure 3.**
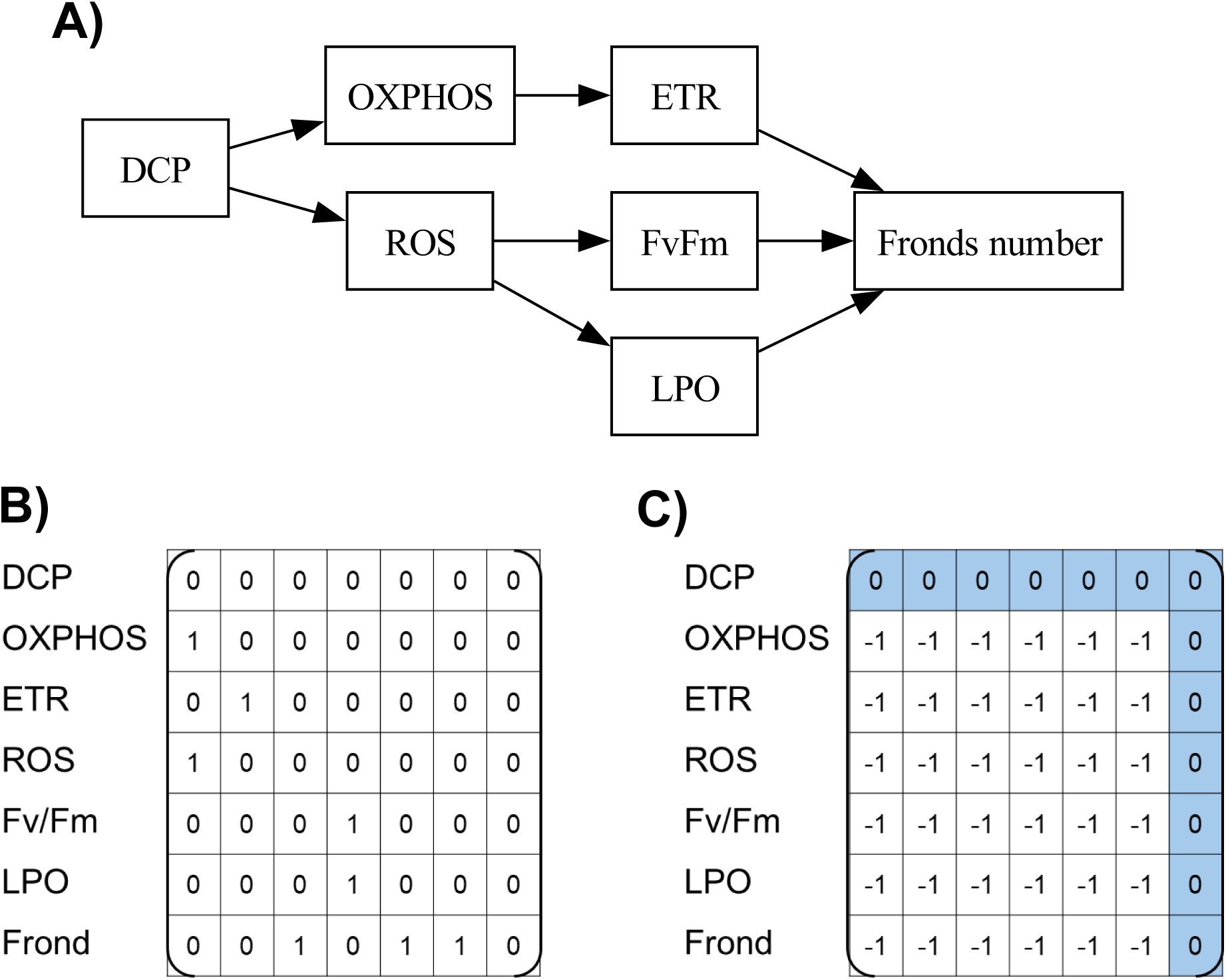
Expert-curated ground-truth graph for 3,5-dichlorophenol (DCP) dataset from Moe et al. (2020). (A) Directed acyclic graph (DAG). (B) Binary adjacent matrix of the ground truth graph. (C) Prior knowledge constraint matrix supplied to DirectLiNGAM. Zero denotes no directed edge, while −1 indicates the edge unconstrained.

We compared each inferred edge across a total of 250,000 iterations with the expert-curated ground-truth graph (Moe et al., 2020) (Figure 3A, 4B) and then calculated arrowheads precision (AHP) and arrowheads recall (AHR) (Zanga et al. 2022): AHP = TP / (TP + FP), AHR = TP / (TP + FN). TP (true positive) denotes edges whose presence and direction match the ground truth, FP (false positive) denotes an inferred edge whose direction or presence is wrong, and FN (false negative) denotes true causal edges that were missed in inferred graphs. For each ordered pair of variables, we also calculated the edge probability—the proportion of iterations in which a given edge appeared—and, analogously, the graph probability for every distinct DAG inferred.

## 3. Results and Discussion

### 3.1. Case study 1: bivariate dose-response relationship

DirectLiNGAM correctly identified concentration as the parent variable with bootstrap probabilities ≥ 96% for dry weight, offspring, and cell viability (fluorescence) (Figures 4A–E). Despite the high statistical reliability, diagnostic hypothesis tests showed that these relationships did not meet at least one of the DirectLiNGAM assumptions. For dry weight and fluorescence, assumptions of linearity and independence of error terms were not supported, while non-Gaussian distribution was supported by the Shapiro-Wilk test (Table 3). For offspring, HSIC test indicated dependence between error terms, although assumptions of linearity and non-Gaussianity were supported. HSIC test indicated violations of independence for all three pairwise relationships, which was consistent with the high F-correlation values (Table 3). These results indicate that DirectLiNGAM can infer the correct causal direction of ecotoxicology data with high statistical reliability, even when some assumptions are violated.

**Figure 4.**
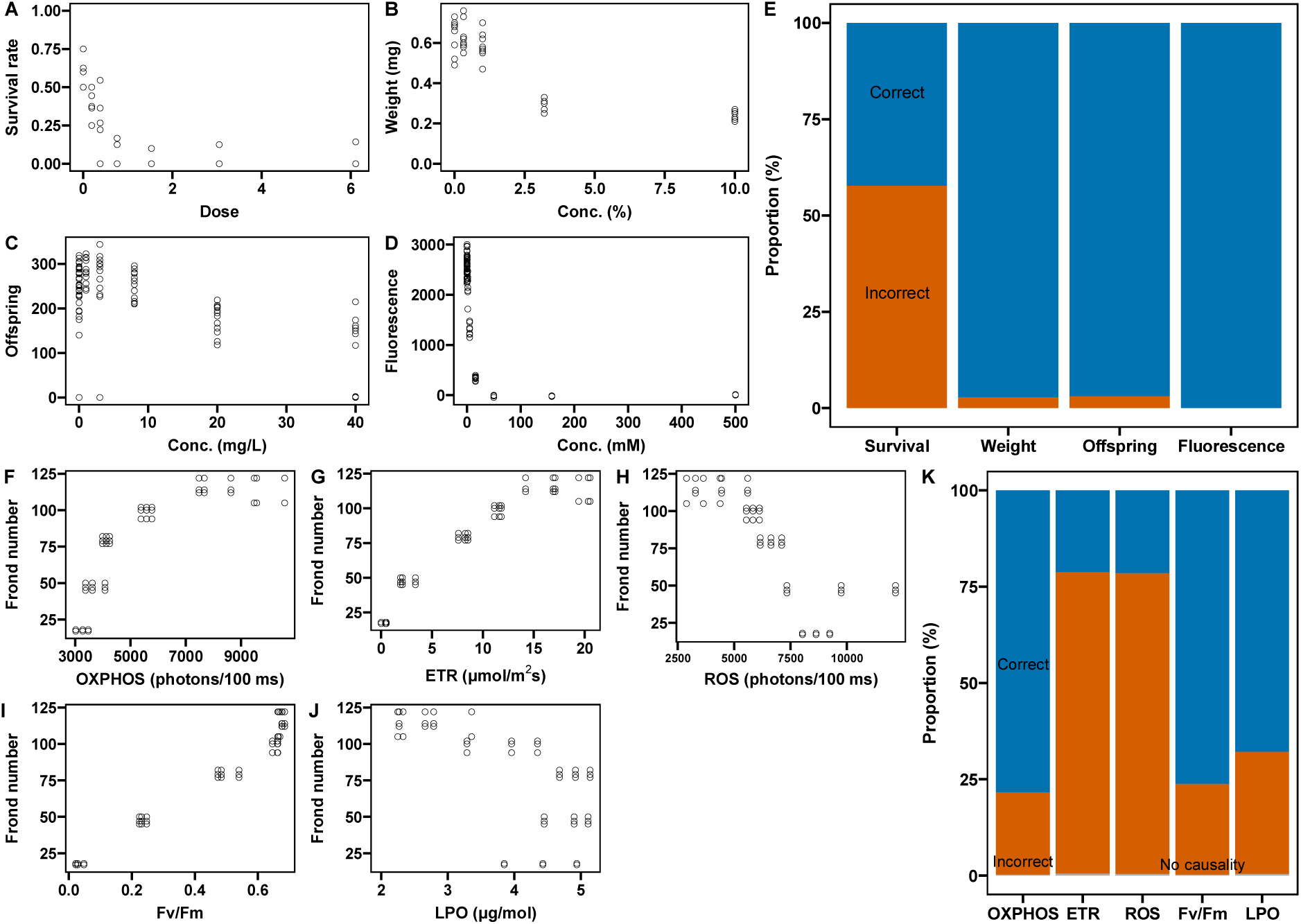
Performance of DirectLiNGAM on bivariate ecotoxicology datasets with known causal direction. Upper and lower panels represent (A–E) dose-response (F–K) and response-response relationships, respectively. (A–D) Scatter plots show the concentration (parent variable) versus the biological response (child variable). (F–J) Scatter plots show five key events (parent variable) plotted against frond number (child variable). (E and K) Bootstrap results expressed as the proportion of bootstrap iterations in which DirectLiNGAM recovered the correct direction (blue), the incorrect direction (orange), or reported no causal link (grey).

**Table 3.** Violations of DirectLiNGAM assumptions in dose-response relationships.

| Relationship | RESET<br>H0: linearity | Shapiro-Wilk test<br>H0: Gaussian† | HSIC test<br>H0: Independent errors | F-correlation |
| --- | --- | --- | --- | --- |
| Concentration → Survival | $p < 0.001$ * | $p = 0.146$ * | $p < 0.001$ * | 0.691 |
| Concentration → Weight | $p < 0.001$ * | $p = 0.007$ | $p < 0.001$ * | 0.754 |
| Concentration → Offspring | $p = 0.371$ | $p < 0.001$ | $p = 0.028$ * | 0.431 |
| Concentration → Fluorescence | $p < 0.001$ * | $p < 0.001$ | $p < 0.001$ * | 0.488 |

Asterisks indicate results that are inconsistent with DirectLiNGAM assumptions.
† For visual inspection of residual distributions, see Figure S3.

**Table 4.** Violations of DirectLiNGAM assumptions in response-response relationships.

| Relationship | RESET<br>H0: linearity | Shapiro-Wilk test<br>H0: Gaussian | HSIC test<br>H0: Independent errors | F-correlation |
| --- | --- | --- | --- | --- |
| OXPHOS -> Frond | $p < 0.001$ * | $p < 0.001$ | $p < 0.001$ * | 0.826 |
| | ($<0.001$–$<0.001$) | ($<0.001$–$<0.001$) | ($<0.001$–$<0.001$) | (0.279–0.908) |
| ETR -> Frond | $p < 0.001$ * | $p < 0.001$ | $p < 0.001$ * | 0.878 |
| | ($<0.001$–$<0.001$) | ($<0.001$–0.003) | ($<0.001$–$<0.001$) | (0.594–0.941) |
| ROS -> Frond | $p < 0.001$ * | $p < 0.001$ | $p < 0.001$ * | 0.819 |
| | ($<0.001$–0.674) | ($<0.001$–0.036) | ($<0.001$–$<0.001$) | (0.372–0.920) |
| Fv/Fm -> Frond | $p = 0.016$ * | $p < 0.001$ | $p < 0.001$ * | 0.894 |
| | (0.001–0.180) | ($<0.001$–0.004) | ($<0.001$–$<0.001$) | (0.449–0.934) |
| LPO -> Frond | $p < 0.233$ | $p < 0.001$ | $p < 0.001$ * | 0.703 |
| | (0.001–0.996) | ($<0.001$–0.001) | ($<0.001$–$<0.001$) | (0.377–0.902) |

Asterisks indicate results that are inconsistent with DirectLiNGAM assumptions. The median p value or F correlation from 25,000 iterations is shown in the top row of each cell, with the range presented in parentheses below.

In contrast, the bootstrap probability for the correct direction of concentration-survival rate relationship was only 42% (Figure 4), indicating that DirectLiNGAM could not clearly identify the causal direction for this pair. The RESET indicated significant non-linearity (*p* < 0.001) for this relationship, and the Shapiro–Wilk test failed to reject the null hypothesis of Gaussian distribution (*p* = 0.146). In addition, the HSIC test indicated dependence between the error terms (*p* < 0.001). These results showed that all LiNGAM assumptions, except for acyclicity, were violated for the concentration-survival rate relationship, which may explain the low bootstrap support and the algorithm’s inability to identify a single causal direction for this pair.

Since ecotoxicology research sometimes involves data transformation to meet statistical assumptions, we tested DirectLiNGAM after the survival rate was transformed using a logit function, and concentration was log-transformed, which is a common approach in ecotoxicology. Although this transformation increased the bootstrap probabilities from 42% to 60% (Figure S2), as with the untransformed data, all three tests still indicated violations of DirectLiNGAM assumptions.

### 3.2. Case study 2: bivariate response-response relationship

For the response–response pairs, the performance of DirectLiNGAM depended on KEs: the pairs between frond number and OXPHOS, Fv/Fm, and LPO showed high bootstrap support (>70%) for the correct causal direction. In contrast, ETR and ROS showed much lower bootstrap support (23% and 28%, respectively) (Figure 3). Although across all five variable pairs, no-causality outputs were rare (< 1 % in every case), both causal directions appeared with > 5% bootstrap probabilities, which indicates that neither direction could be fully excluded.

The reason why DirectLiNGAM could not determine a single causal direction reliably for response-response relationships remains unclear; however, one likely factor is the limited number of replicates per concentration (i.e., *n* = 3). Diagnostic hypothesis tests showed that all variable pairs satisfied the non-Gaussian assumption yet violated the independence and linearity assumptions, except for LPO. Because these assumption violations were similar to those observed in the dose-response case studies, they are unlikely to explain the reduced reliability for response-response relationships. To further explore this issue, we examined the impact of replicate numbers on causal direction estimates. When the number of replicates per concentration was reduced from 8 to 2 for the concentration-weight and concentration-fluorescence relationships, the average bootstrap probability for the correct direction decreased from > 95% to 74.3 ± 2.7% (weight) and to 93.5 ± 0.9% (fluorescence) (Figure S4), suggesting that even well-supported causal pairs can lose statistical reliability for direction under reduced replication. This result suggests that a sufficient number of replicates is required for statistical reliability in causal direction inference. Despite the limited replication in the DCP dataset, we used it as a multivariate case study (section 3.3) because alternative datasets with known causal structure were not available.

### 3.3. Case study 3: multivariate relationship

Without any prior knowledge, the ensemble of inferred DAGs showed substantial iteration-to-iteration variability. The single most common graph appeared in only 0.04% of runs (Figure 5A); the next three graphs occurred at 0.03%, 0.02%, and 0.02%, respectively. In addition, these DAGs rarely shared causal edges (Table S2 and Figure S5); no edges were commonly observed in all ten most frequent DAGs. These very low probabilities indicate high structural uncertainty. In addition, these DAGs contained mechanistically implausible edges, such as from Fv/Fm to DCP concentration and from ETR to DCP, suggesting the limitations of an unconstrained search. Consequently, even ten most frequent DAGs were far from the expert ground truth, with AHP of 0.08–0.23 and AHR of 0.13–0.38 (Table S2).

**Figure 5.**
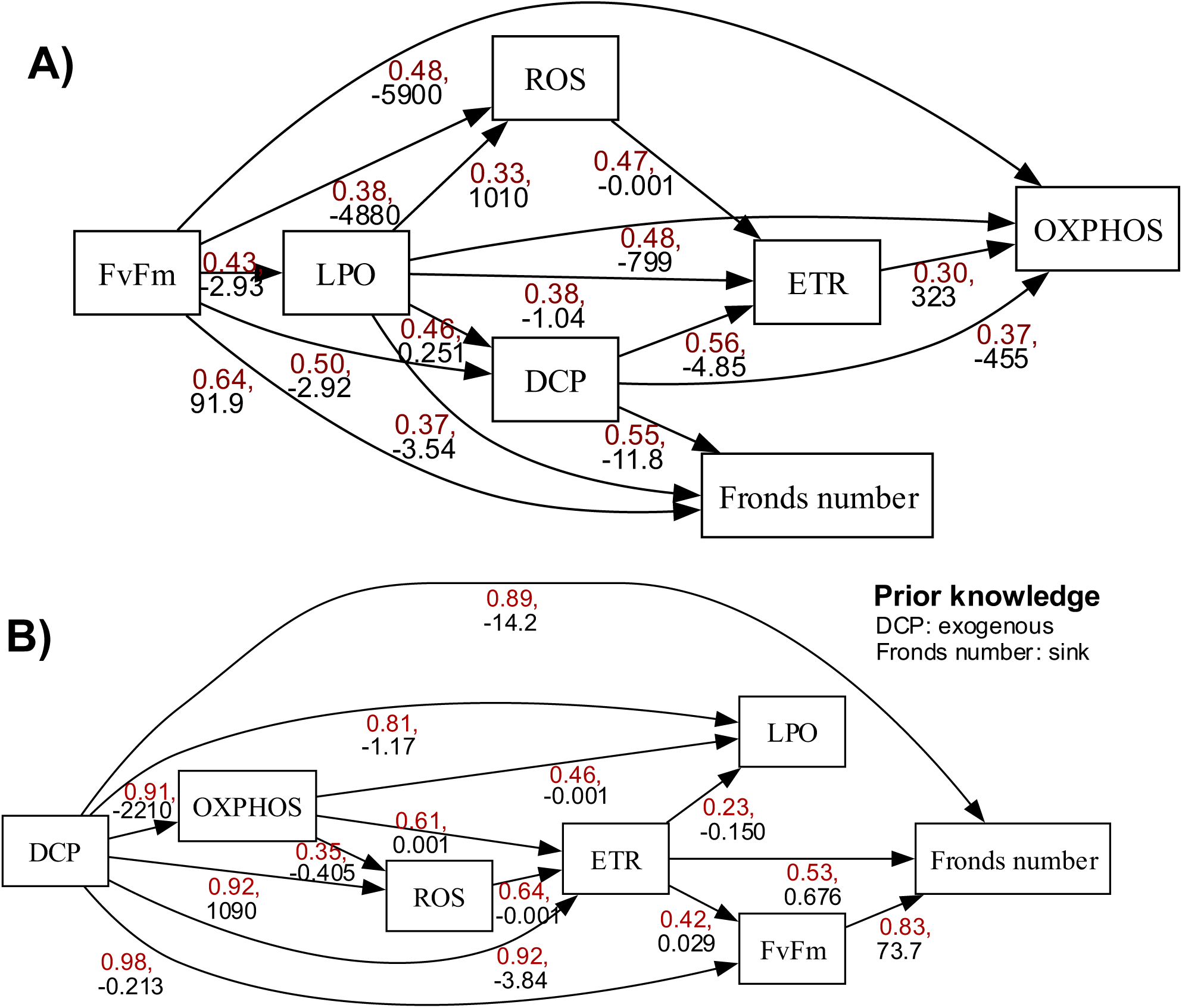
Most frequent causal graph for 3,5-dichlorophenol (DCP) dataset inferred by DirectLiNGAM (A) without and (B) with prior knowledge. For each arrow, the upper red value is the edge probability—the proportion iterations (out of 250,000) in which the edge appeared, while the lower black value represents the mean standardized direct causal effect across runs for the most frequent DAG.

Adding minimal prior information substantially improved the outcome. By declaring DCP concentration as exogenous and frond number as a sink variable, the most frequent graph (Figure 5B) more closely resembled the expert ground truth, with AHP of 0.36 and AHR of 0.63. Although the absolute probability of any single DAG remained low (Table S3), the inferred graphs now shared consistent causal motifs. For example, edges from DCP to OXPHOS, from DCP to ROS, and from OXPHOS to LPO were observed in all ten most frequent DAGs (Table S3 and Figure S6). The AHP and AHR were 0.31–0.40 and 0.50–0.75, respectively, for ten most frequent DAGs. Furthermore, OXPHOS and ROS located in the upstream of graph, which is consistent with the expert ground truth with these variables as MIE, although the estimated contributions of paths through these variables were not high (see next paragraph).

To compare pathway contributions to the AO (i.e., reduced frond number), we calculated path probabilities and total causal effects from DCP to frond number (Table 5) using the most-frequent, prior knowledge-constrained DAG. Ground-truth paths, such as “ DCP → OXPHOS → ETR → frond number” and “ DCP → ROS → LPO → frond number”, were observed with non-negligible probabilities and effects. However, the direct path of DCP → frond number exhibited the largest probability and large negative effect (−21.12), suggesting that unmeasured KEs may still be missing from the current model.

**Table 5.** Estimated probabilities and total causal effects for paths starting from 3,5-dichlorophenol (DCP) to frond number.

| Path | Probability | Causal effect |
| --- | --- | --- |
| DCP→Frond | 0.89 | -19.00 |
| DCP→Fv/Fm→Frond | 0.82 | -21.12 |
| DCP→ETR→Frond | 0.50 | -3.96 |
| DCP→OXPHOS→Frond | 0.48 | 3.86 |
| DCP→LPO→Frond | 0.43 | -0.75 |
| DCP→ROS→Frond | 0.37 | -1.03 |
| DCP→ETR→Fv/Fm→Frond | 0.34 | -4.74 |
| **DCP→OXPHOS→ETR→Frond** | **0.33** | **-2.09** |
| DCP→ROS→ETR→Frond | 0.33 | -0.68 |
| DCP→LPO→Fv/Fm→Frond | 0.28 | 1.08 |
| DCP→OXPHOS→LPO→Frond | 0.23 | -1.90 |
| DCP→OXPHOS→ETR→Fv/Fm→Frond | 0.23 | -3.25 |
| ... | ... | ... |
| **DCP→ROS→LPO→Frond** | **0.13** | **-0.84** |
| ... | ... | ... |
| **DCP→ROS→Fv/Fm→Frond** | **0.001** | **2.25** |

Paths with the highest probabilities are presented. Paths that are part of the expert ground-truth network are shown in bold red.

### 3.4. Implications and further research recommendations

This study demonstrates the strategic value of SCD as a novel tool for AOP development and evaluation. Although algorithms such as DirectLiNGAM impose several assumptions, our bivariate examples show that SCD can still infer the correct causal direction of ecotoxicology data even when some assumptions are violated. The multivariate case study also illustrates that DirectLiNGAM can partially infer biologically plausible causal structure in a data-driven manner.

SCD can contribute to AOP science in two ways. First, it can be useful in developing unknown AOP in a data-driven manner. In the DCP multivariate case study, DirectLiNGAM correctly located OXPHOS inhibition and ROS elevation upstream, identifying them as candidate MIEs. Pinpointing MIEs is critical for defining an AOP’s taxonomic applicability domain and for designing relevant in vitro assays (Russom et al. 2014; Fay et al. 2017). Although omics-integrated SCD was beyond this work’s scope, such a combination holds great promise. Nevertheless, the high dimensionality of omics datasets, often comprising hundreds to thousands of variables, exceeds the capacity of conventional SCD algorithms. To address this, it is recommended to apply high-dimensional SCD algorithms (Samuel Wang and Drton 2020) and pre-filtering methods, such as statistical testing and machine learning-based feature selection (Tamura et al. 2024).

Second, SCD provides an objective means to test and refine existing AOP and qAOP networks. AOP networks are typically built by integrating multiple linear AOPs that were developed manually (Knapen et al. 2018), meaning that the graph skeleton of a qAOP often depends on prior knowledge derived from AOP databases or expert judgement. SCD can identify missing links between existing KEs. For example, in the DCP case study, the OXPHOS → ETR → Fv/ m path was detected with high probability (23%) and substantial causal effect (Table 5), which was absent from the original ground-truth graph. This path is biologically plausible, because ATP depletion caused by OXPHOS impairment can suppress photosystem II recovery (Singh et al. 1996) without ROS mediation. Thus, the inclusion of this path may improve the predictive power of qAOP. Although AOP and AOP networks are not intended to fully capture the complexity of biological systems (Villeneuve et al. 2014), causal pathways with strong influence on the AO should be considered for inclusion. This refinement process is fully consistent with recent calls to adopt KE relationships, rather than linear AOP, as the pragmatic unit of AOP development (Foran et al. 2019; Svingen et al. 2021). In addition, by comparing SCD-inferred graphs with the expert-curated drafts, we can identify strongly and weakly supported KE relationships and prioritize follow-up experiments. This approach would serve as a valuable complement to existing methods, where the reliability of each KE relationship is assessed through expert judgement (Becker et al. 2015).

We selected simple examples as the initial applications of SCD to AOP development. However, there are many potential directions to expand the use of SCD. First, it is possible to relax the assumption of acyclicity, commonly required by many SCD algorithms, including LiNGAM, as biological systems often contain feedback loops. Indeed, AOP and AOP networks allow cyclic structures (OECD 2018; Pollesch et al. 2019), although bidirectional edges between two KEs (i.e., two headed arrows) are not allowed (OECD 2018). To accommodate such biological feedback, cycle-tolerant SCD algorithms could be explored, such as an extension of LiNGAM (Lacerda et al. 2008) and additive noise models with feedback loops (Mooij et al. 2011). Second, it is important to consider SCD algorithms that account for hidden common causes. While chemical concentration or stressor level can be modeled as exogenous, multiple KEs, including MIEs and AOs, may share unmeasured causes. In such cases, causal inference using standard SCD methods may yield biased or uncertain results. Other SCD approaches could be useful, such as constraint-based approaches that allow for hidden variables (e.g., FCI) and an extension of LiNGAM that tolerate hidden common causes (Maeda and Shimizu 2020). Third, incorporating more detailed prior knowledge may further enhance SCD performance. Although only DCP and frond number were treated as the exogenous variable and sink variable, respectively, as a minimal form of prior knowledge in the DCP multivariate case example, additional constraints could be applied. For example, AOPs typically span multiple levels of biological organization, where effects at the molecular and cellular levels generally precede those at the organ and individual levels. This hierarchical understanding can be encoded as prior knowledge, thereby narrowing the search space and improving the accuracy of SCD. Lastly, the use of time-course data may improve causal inference by leveraging temporal order, since causes must precede their effects. Several SCD methods for time series exist, such as Granger causality test and VARLiNGAM (Hyvarinen et al. 2010), and have been successfully applied in molecular biology and gene regulatory network inference (Heerah et al. 2021; Schlamp et al. 2021). Although time-course data remain relatively scarce in ecotoxicology, particularly for molecular and cellular-level endpoints (Ankley and Villeneuve 2015; Schüttler et al. 2019) likely due to the need for destructive sampling, such data could greatly enhance the applicability and accuracy of SCD in future studies.

In addition to advancing SCD methodologies, the establishment of non-destructive sampling techniques would be beneficial for SCD-driven AOP development. Examples of such techniques are phenotypic observation (Michaelis et al. 2025), behavioral tracking (Tamura et al. 2024), and waterborne metabolite or RNA analysis (Song et al. 2017; Hiki et al. 2023). These methods enable the collection of datasets without missing values, and their ability to support repeated measurements facilitates the generation of time-course data, thereby enhancing the robustness and efficiency of causal inference.

Finally, we highlight key considerations when applying SCD to AOP development. As reflected by the low AHP and high AHR in the DCP multivariate case example, the inferred graphs may contain false positives. Some of these may represent biologically or toxicologically unrealistic paths. In such cases, these pathways should be filtered out using additional sources, such as expert knowledge. SCD is not a silver bullet and its combination with complementary tools, such as LLM (Takayama et al. 2025) and machine learning (Tamura et al. 2024), would enhance overall reliability. In addition, as shown by low DAG bootstrap probabilities in the DCP cases, ecotoxicological studies often involve small sample sizes, leading to greater uncertainty. We encourage future applications of SCD in ecotoxicology to incorporate larger numbers of biological replicates to improve robustness and reproducibility of causal inference.

## ASSOCIATED CONTENT

### Supporting Information

Performance of analysis methods for block-wise missing data (Table S1); Ten most frequent DAGs inferred by DirectLiNGAM without prior knowledge (Table S2); Ten most frequent DAGs inferred by DirectLiNGAM with prior knowledge (Table S3); Example of simulated block-wise missing data (Figure S1); Performance of DirectLiNGAM on bivariate ecotoxicology dataset with logit and logarithmic transformation (Figure S2); Observed distribution of residuals (Figure S3); Bootstrap probability across different numbers of replicates (Figure S4); Ten most frequent DAGs inferred by DirectLiNGAM without prior knowledge (Figure S5); Ten most frequent DAGs inferred by DirectLiNGAM with prior knowledge (Figure S6) (PDF)

## AUTHOR INFORMATION

### Author contributions

Kyoshiro Hiki: Conceptualization, Methodology, Investigation, Formal analysis, Writing – original draft, review & editing. Thong Pham: Conceptualization, Methodology, Formal analysis, Validation, Writing – review & editing. Michio Yamamoto: Methodology, Formal analysis, Writing –review & editing. Takehiko I Hayashi: Conceptualization, Methodology, Formal analysis, Funding acquisition, Writing – original draft, review & editing. Shohei Shimizu: Conceptualization, Methodology, Project administration, Funding acquisition, Writing –review & editing.

### Notes

The authors declare no conflicts of interest. The opinions expressed in this manuscript do not necessarily represent the official views of the authors’ affiliation.

## ACKNOWLEDGMENTS

This study was financially supported by JST, CREST (Grant Number: JPMJCR22D2).

