## Supporting Information for "Statistical Causal Discovery in Developing and Refining Adverse Outcome Pathway (AOP)"

### Table of contents


| | | Content | Page |
|---|---|---|---|
| **Text** | Text 1 | Comparison of imputation / estimation approaches for block-wise missing data | S3 |
| **Tables** | Table S1 | Performance of analysis methods for block-wise missing data | S5 |
| | Table S2 | Ten most frequent DAGs inferred by DirectLiNGAM without prior knowledge | S6 |
| | Table S3 | Ten most frequent DAGs inferred by DirectLiNGAM with prior knowledge | S7 |
| **Figures** | Figure S1 | Example of simulated block-wise missing data | S3 |
| | Figure S2 | Performance of DirectLiNGAM on bivariate ecotoxicology dataset with logit and logarithmic transformation | S8 |
| | Figure S3 | Observed distribution of residuals | S9 |
| | Figure S4 | Bootstrap probability across different numbers of replicates | S10 |
| | Figure S5 | Ten most frequent DAGs inferred by DirectLiNGAM without prior knowledge | S11 |
| | Figure S6 | Ten most frequent DAGs inferred by DirectLiNGAM with prior knowledge | S12 |

### Text S1: Comparison of imputation / estimation approaches for block-wise missing data

Using simulated data, we compared five methods for handling common ecotoxicology data in which each endpoint is measured in a separate replicate—i.e., the block-wise missing pattern where only concentration ($Z$) and one biological endpoint ($X_1$, $X_2$, or $Y$) are recorded per individual. In this preliminary investigation, the relationships between variables were defined as follows:

$$X_1 = a_1 + b_1 Z + e_1 \qquad e_1 \sim N(0, \sigma_1^2)$$

$$X_2 = a_2 + b_2 Z + e_2 \qquad e_2 \sim N(0, \sigma_2^2)$$

$$Y = g_0 + g_1 X_1 + g_2 X_2 + e_y \qquad e_y \sim N(0, \sigma_y^2)$$

The coefficients were defined as follows: $a_1 = 2$, $b_1 = 1$, $a_2 = 1$, and $b_2 = 0.5$; $g_1$ and $g_2$ representing causal effects, were set to 1.2 and 0.8, respectively. The variances were set as follows: $\sigma_1 = 1$, $\sigma_2 = 1$, and $\sigma_y = 0.1$. Concentration Z was discrete variable and took 10 distinct levels (0–9). For each level, fifteen replicates were generated with three complete variables ($X_1$, $X_2$, and $Y$) (Figure S1A), and two of the three variables were then deleted, yielding a table in which each endpoint column contained two-thirds missing values (Figure S1B).

**A) Full data**

| Z (Concentration) | X1 | X2 | Y |
|---|---|---|---|
| 0 | 1.7 | 1.9 | 5.7 |
| 0 | 3.0 | 2.1 | 5.2 |
| 0 | 3.4 | 0.1 | 2.5 |
| 0 | 1.6 | 0.5 | 3.9 |
| 0 | 1.3 | 1.5 | 4.6 |
| 0 | 1.9 | 2.5 | 5.1 |
| 0 | 1.1 | 0.4 | 3.8 |
| 0 | 2.5 | -0.1 | 4.9 |
| 0 | 1.5 | 1.7 | 2.0 |
| 0 | 2.0 | 0.9 | 3.1 |
| 0 | 2.3 | 0.8 | 4.1 |
| 0 | 1.8 | 1.0 | 3.7 |
| 0 | 2.1 | -0.2 | 2.9 |
| 0 | 1.5 | 0.3 | 3.2 |
| 0 | 1.9 | 1.2 | 3.5 |
| 1 | 3.0 | 2.7 | 5.7 |
| … | … | … | … |
| 9 | 12.1 | 6.4 | 18 |

**B) Data with missing values**

| Z (Concentration) | X1 | X2 | Y |
|---|---|---|---|
| 0 | 1.7 | NA | NA |
| 0 | 3.0 | NA | NA |
| 0 | 3.4 | NA | NA |
| 0 | 1.6 | NA | NA |
| 0 | 1.3 | NA | NA |
| 0 | NA | 2.5 | NA |
| 0 | NA | 0.4 | NA |
| 0 | NA | -0.1 | NA |
| 0 | NA | 1.7 | NA |
| 0 | NA | 0.9 | NA |
| 0 | NA | NA | 4.1 |
| 0 | NA | NA | 3.7 |
| 0 | NA | NA | 2.9 |
| 0 | NA | NA | 3.2 |
| 0 | NA | NA | 3.5 |
| 1 | 3.0 | NA | NA |
| … | … | … | … |
| 9 | NA | NA | 18 |

Figure S1. Example of simulated data for comparison of analysis methods. After generation of a full data set (A), two variables were deleted within each $Z$ level, to mimic the common missing pattern in ecotoxicology.

The simulated data with missing values were subjected to five methods: i) MEAN-PICK, ii) RANDOM-PICK, iii) IMPUTE-S, iv) IMPUTE-M, and v) RAW-IV. The estimated direct effects $g_1$ and $g_2$ were compared with their true values (i.e., 1.2 and 0.8). The details of each method are as follows:

- ✓ i) MEAN-PICK replaces each missing value with the mean of the observed values from the same $Z$ level, and then performs ordinary least squares (OLS) regression: $Y \sim X_1 + X_2$.
- ✓ ii) RANDOM-PICK randomly replaces each missing values with only one observed value from the same $Z$ level, performs OLS regression, repeats these steps 100 times, and calculates the mean of regression coefficients.
- ✓ iii) IMPUTE-S replaces missing values by sampling from observed values from the same $Z$ level, and then performs OLS regression.
- ✓ iv) IMPUTE-M similarly replaces missing values by sampling from observed values from the same $Z$ level, performs OLS regression, repeats these steps 100 times, and calculates the mean of regression coefficients.
- ✓ v) RAW-IV does not impute missing values, but estimates $g_1$ and $g_2$ directly via $Z$-instrumented instrumental variables (IV): Cov $(Y, Z)$/Cov $(X, Z)$

For all the methods, we generated 100 datasets and calculated standard deviations (SD) of the estimated coefficients across 100 iterations and root mean squared error (RMSE = $\sqrt{\frac{1}{N}\sum_{k=1}^{N}(\hat{g} - g)^2}$) as performance metrics.

As a result, IMPUTE-M achieved the lowest RMSE values for both direct causal effects $g_1$ and $g_2$ (Table S1), followed by RANDOM-PICK, IMPUTE-S, MEAN-PICK, and RAW-IV. Furthermore, the mean causal effects estimated by IMPUTE-M were close to the true values. These preliminary results suggest that IMPUTE-M (i.e., multiple imputation) is a practical and statistically reliable approach for handling block-wise missing ecotoxicology data.

Table S1. Performance of five analysis methods to estimate direct causal effects $g_1$ and $g_2$.

| | $g_1$ (True value: 1.20) | | | $g_2$ (True value: 0.80) | | |
|---|---|---|---|---|---|---|
| | Mean | SD | RMSE | Mean | SD | RMSE |
| i) MEAN-PICK | 1.28 | 0.39 | 0.39 | 0.59 | 0.71 | 0.74 |
| ii) RANDOM-PICK | 1.20 | 0.12 | 0.12 | 0.59 | 0.19 | 0.28 |
| iii) IMPUTE-S | 1.17 | 0.13 | 0.14 | 0.59 | 0.20 | 0.29 |
| iv) IMPUTE-M | 1.17 | 0.11 | **0.11** | 0.60 | 0.17 | **0.27** |
| v) RAW-IV | 1.61 | 0.11 | 0.43 | 3.15 | 0.33 | 2.37 |

Bold indicates the lowest RMSE values.

Table S2. Ten most frequent directed acyclic graphs (DAGs) inferred by DirectLiNGAM without prior knowledge.

| | Graph* | Frequency | True positive | False positive | False negative | Arrowheads precision | Arrowheads recall |
|---|---|---|---|---|---|---|---|
| 1 | 0000110 1010110 1001010 0000110 0000000 0000100 1000110 | 0.04% | 3 | 12 | 5 | 0.200 | 0.375 |
| 2 | 0010100 1010110 0000100 0010000 0000000 0010100 1010100 | 0.03% | 3 | 10 | 5 | 0.231 | 0.375 |
| 3 | 0100010 0000000 1001010 1000010 1000011 0100000 1110010 | 0.02% | 3 | 12 | 5 | 0.200 | 0.375 |
| 4 | 0100000 0000000 1101000 0100000 1000001 1100001 1101000 | 0.02% | 1 | 12 | 7 | 0.077 | 0.125 |
| 5 | 0010100 1010110 0000100 0010000 0000000 0010100 1000100 | 0.02% | 2 | 10 | 6 | 0.167 | 0.250 |
| 6 | 0100110 0000110 0100100 0100100 0000000 0000100 1000110 | 0.02% | 3 | 10 | 5 | 0.231 | 0.375 |
| 7 | 0000110 1010110 1001110 0000110 0000000 0000100 1010100 | 0.02% | 3 | 13 | 5 | 0.188 | 0.375 |
| 8 | 0010100 1010110 0000101 0010000 0000000 0010100 0000100 | 0.02% | 2 | 10 | 6 | 0.167 | 0.250 |
| 9 | 0010100 1010110 0000100 0010000 0000000 0010100 1000110 | 0.02% | 3 | 10 | 5 | 0.231 | 0.375 |
| 10 | 0100000 0000000 1101000 1100000 1010000 1110000 1010000 | 0.02% | 3 | 10 | 5 | 0.231 | 0.375 |

*: An adjacency matrix is expressed as character strings. The first segment lists edges directed to DCP concentration from all other variables.

See Figure S5 for graphical representations.

Table S3. Ten most frequent directed acyclic graphs (DAGs) inferred by DirectLiNGAM with prior knowledge.

| | Graph* | Frequency | True positive | False positive | False negative | Arrowheads precision | Arrowheads recall |
|---|---|---|---|---|---|---|---|
| 1 | 0000000 1000000 1101000 1100000 1010000 1110000 1010100 | 0.06% | 5 | 9 | 3 | 0.357 | 0.625 |
| 2 | 0000000 1000000 1101000 1100000 1010000 1110000 1110100 | 0.05% | 5 | 10 | 3 | 0.333 | 0.625 |
| 3 | 0000000 1000000 1101000 1100000 1010000 1100100 1010100 | 0.04% | 5 | 9 | 3 | 0.357 | 0.625 |
| 4 | 0000000 1000000 1101000 1000000 1010000 1110000 1010100 | 0.04% | 5 | 8 | 3 | 0.385 | 0.750 |
| 5 | 0000000 1000000 1101000 1100000 1010000 1110000 1110110 | 0.04% | 6 | 10 | 2 | 0.375 | 0.625 |
| 6 | 0000000 1000000 1101000 1100000 1010000 1110000 1110010 | 0.04% | 5 | 10 | 3 | 0.333 | 0.625 |
| 7 | 0000000 1000000 1101000 1100000 1010010 0100000 1110110 | 0.04% | 6 | 9 | 2 | 0.400 | 0.750 |
| 8 | 0000000 1000000 1101000 1100000 1010010 1100000 1000100 | 0.04% | 4 | 9 | 4 | 0.308 | 0.500 |
| 9 | 0000000 1000000 1101000 1100000 1010010 1100000 1000110 | 0.04% | 5 | 9 | 3 | 0.357 | 0.625 |
| 10 | 0000000 1000000 1101000 1100000 1010010 1110000 1010100 | 0.04% | 5 | 10 | 3 | 0.333 | 0.625 |

*: An adjacency matrix is expressed as character strings. The first segment lists edges directed to DCP concentration from all other variables.

See Figure S6 for graphical representations.

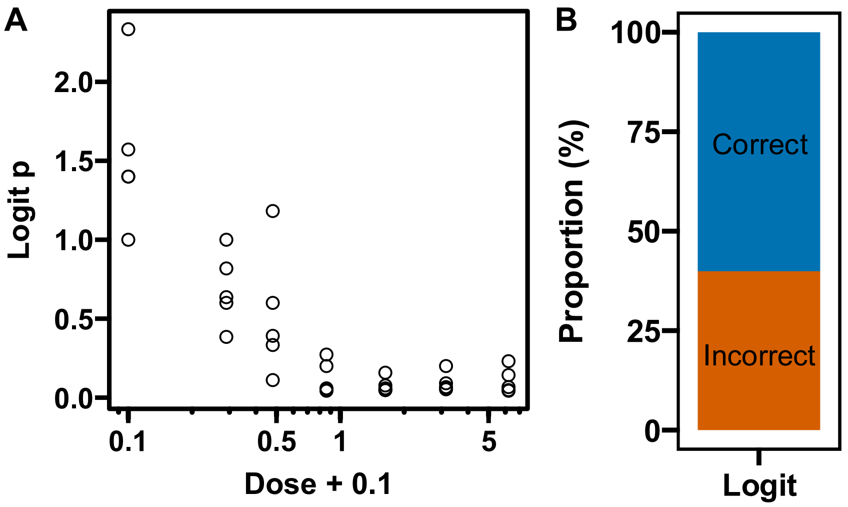


Figure S2. Performance of DirectLiNGAM on bivariate ecotoxicology dataset with logit and logarithmic transformation. (A) Scatter plots show the logarithmic concentration (parent variable) versus the logit function of survival rate (child variable). (B) Bootstrap results expressed as the proportion of iterations in which DirectLiNGAM recovered the correct direction (blue) or the incorrect direction (orange).

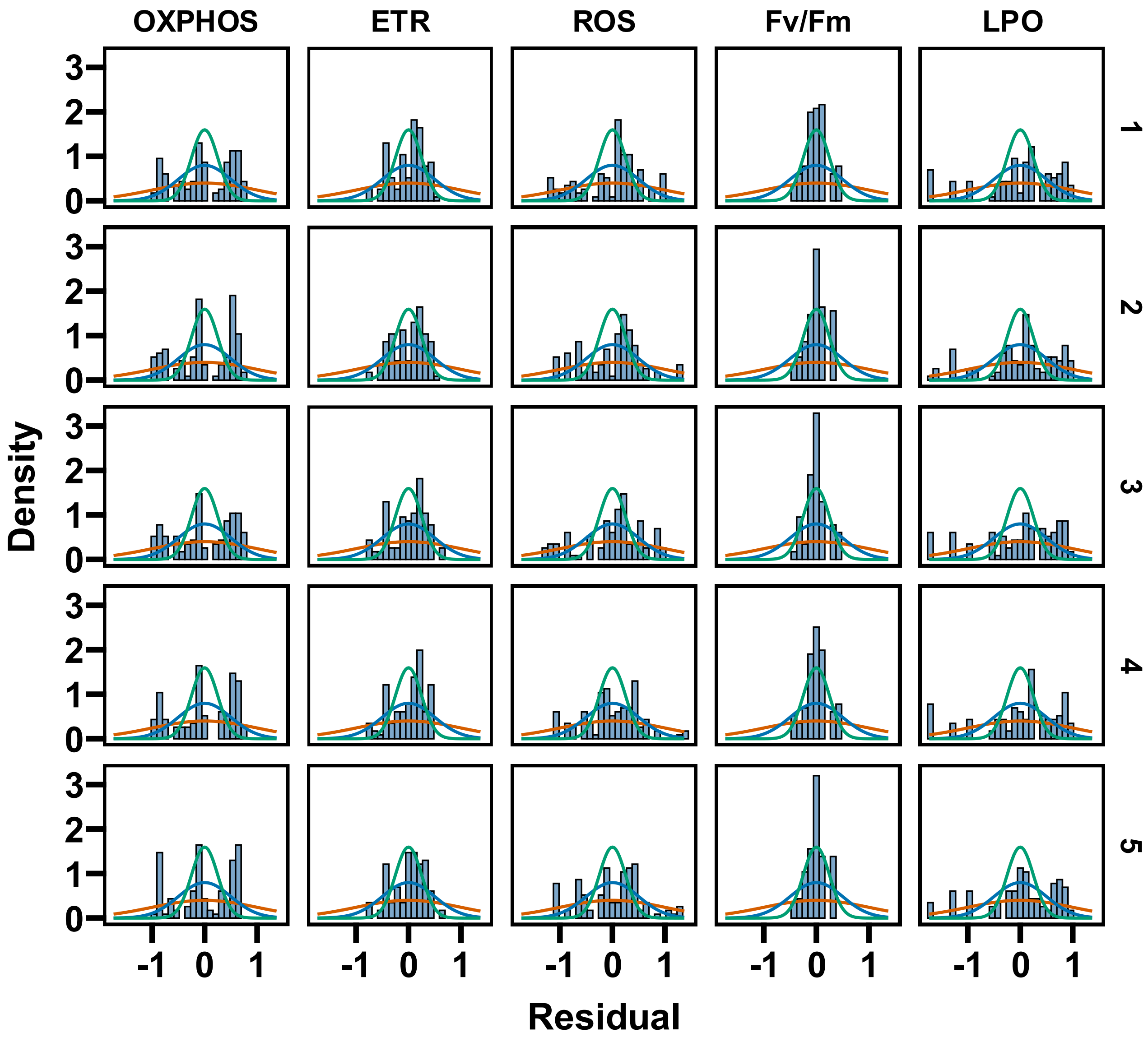


Figure S3. The observed distribution of residuals of linear regressions for upstream endpoint (OXPHOS, ETR, ROS, Fv/Fm, and LPO) against frond number. Orange, blue, and green lines represent normal distributions with standard deviation of 1, 0.5, and 0.25, respectively. Different rows denote data from different iterations.

A)

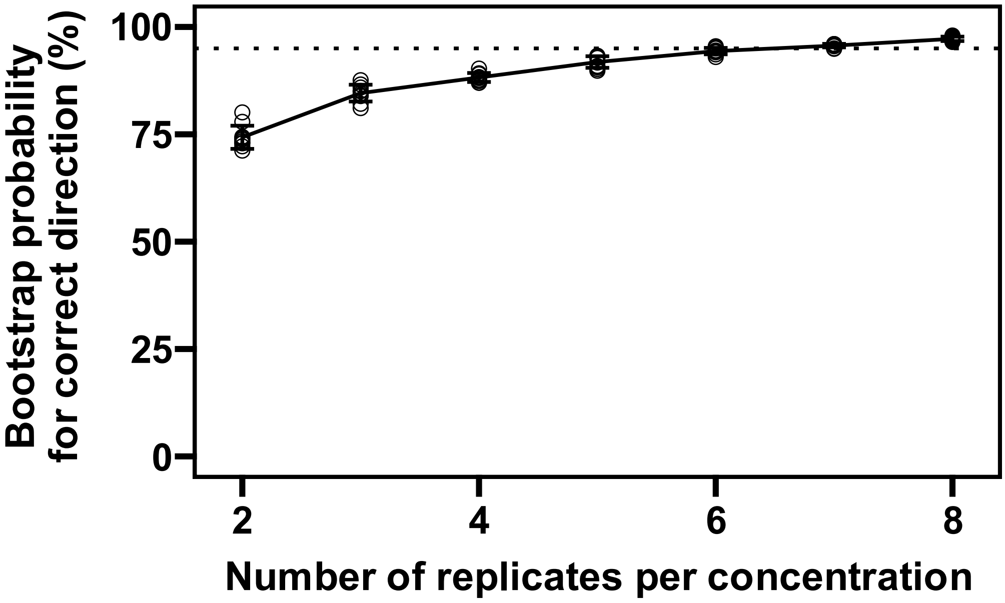


B)

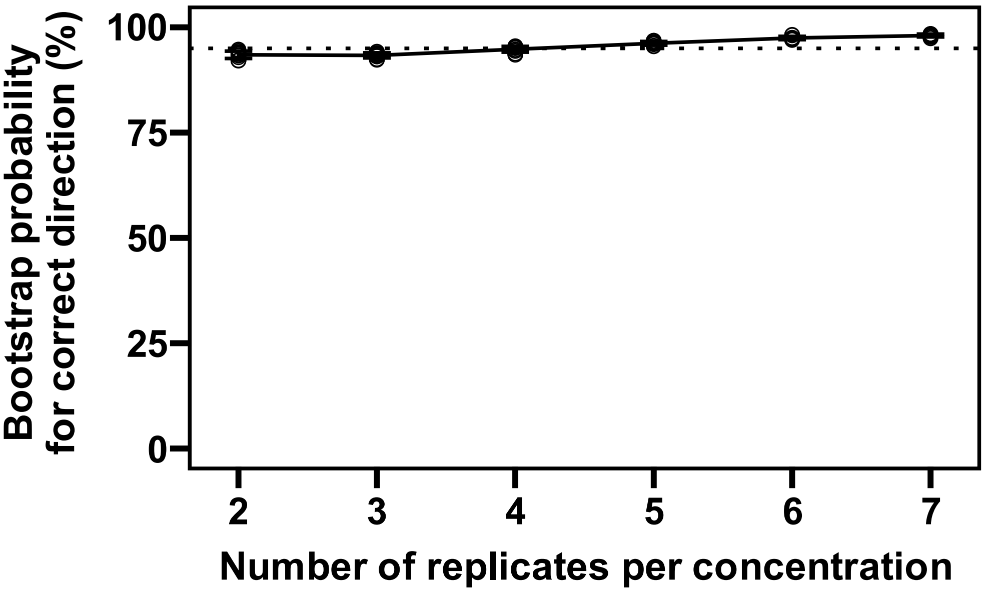


Figure S4. Bootstrap probability of DifrectLiNGAM in inferring correct causal direction for dose-response pairs across different numbers of replicates per concentration. (A) concentration-weight pair, and (B) concentration-fluorescence pair. The bootstrap probabilities were estimated by performing DirectLiNGAM with 1000 bootstrap iterations, while varying the number of replicates per concentration. For each replicate setting, a random selection was performed 10 times. The dotted line indicates a 95% bootstrap probability.

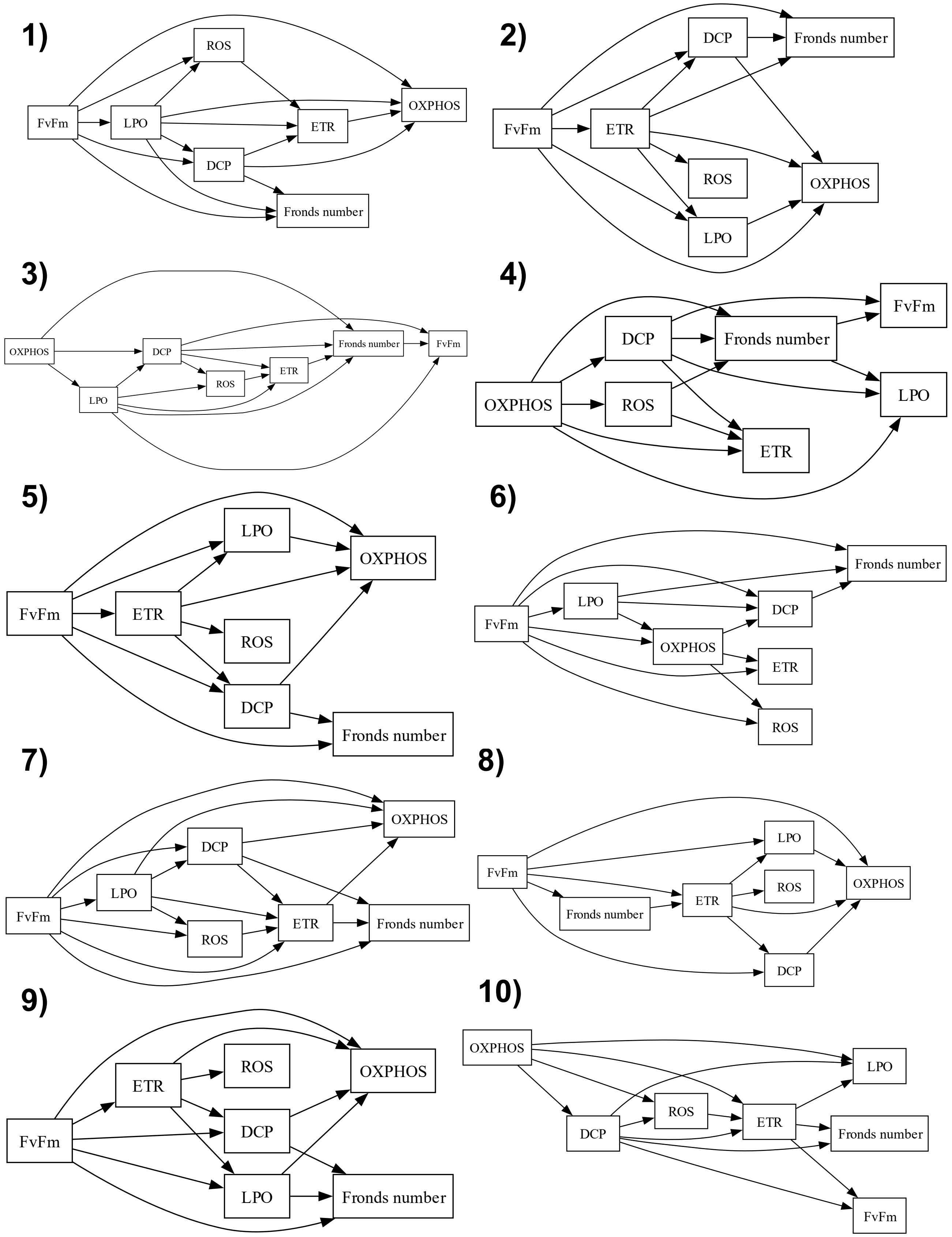


Figure S5. Ten most frequent directed acyclic graphs (DAGs) inferred by DirectLiNGAM without prior knowledge. The numbers correspond to those shown in Table S2.

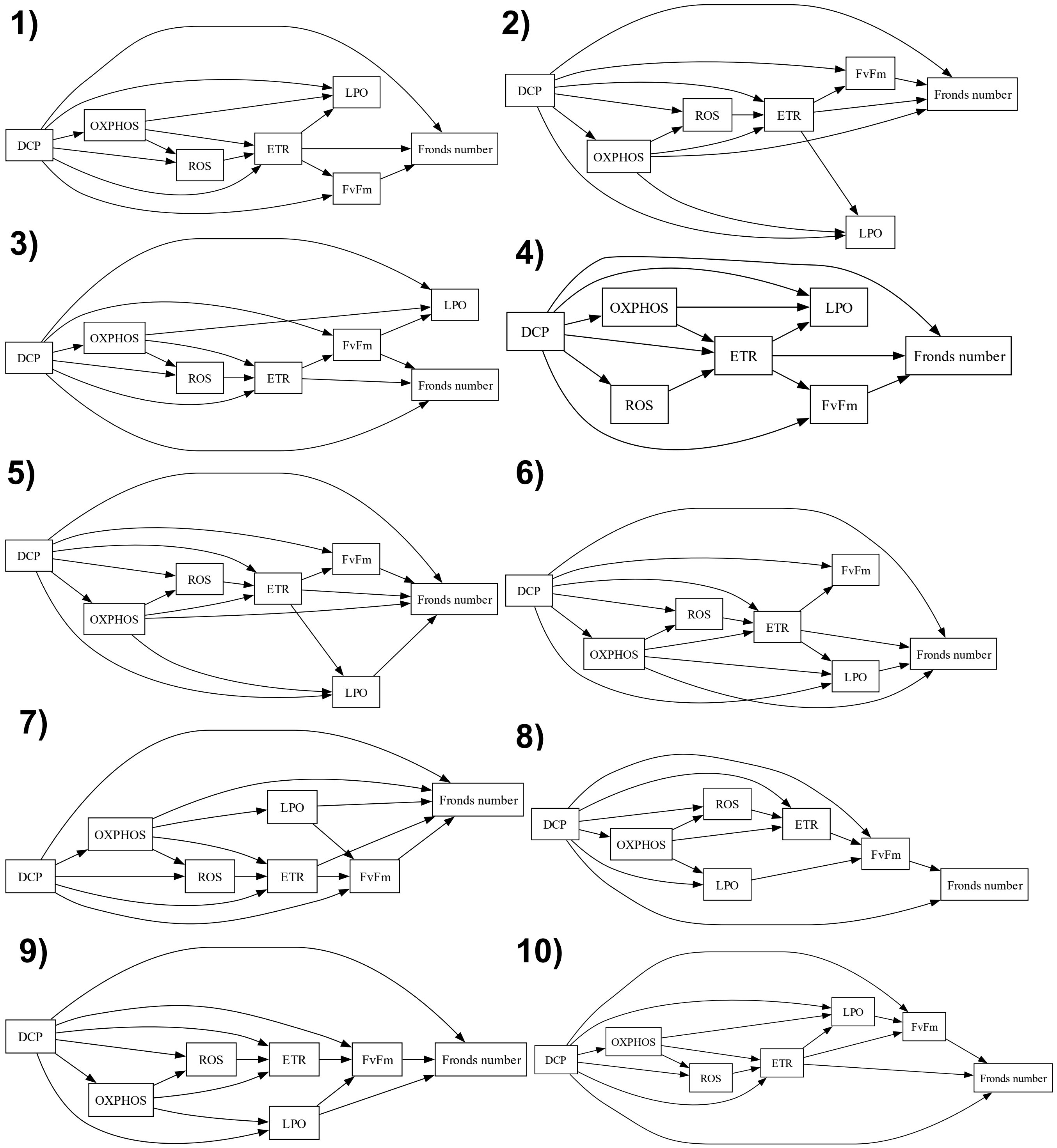


Figure S6. Ten most frequent directed acyclic graphs (DAGs) inferred by DirectLiNGAM with prior knowledge. The numbers correspond to those shown in Table S3.
